# The genetic basis and fitness consequences of sperm midpiece size in deer mice

**DOI:** 10.1101/077826

**Authors:** Heidi S. Fisher, Emily Jacobs-Palmer, Jean-Marc Lassance, Hopi E. Hoekstra

## Abstract

An extraordinary array of reproductive traits vary among species, yet the genetic mechanisms that enable divergence, often over short evolutionary timescales, remain elusive. Here we examine two sister-species of *Peromyscus* mice with divergent mating systems. We find that the promiscuous species produces sperm with longer midpiece than the monogamous species, and midpiece size correlates positively with competitive ability and swimming performance. Using forward genetics, we identify a gene associated with midpiece length: *Prkar1a*, which encodes the R1α regulatory subunit of PKA. R1α localizes to midpiece in *Peromyscus* and is differentially expressed in mature sperm of the two species yet is similarly abundant in the testis. We also show that genetic variation at this locus accurately predicts male reproductive success. Our findings suggest that rapid evolution of reproductive traits can occur through cell type-specific changes to ubiquitously expressed genes and have an important effect on fitness.

---

The remarkable diversity of male reproductive traits observed in nature is often attributed the evolutionary forces of sexual conflict, sperm competition, and sperm precedence^1−4^. However, the genetic mechanisms that enable reproductive traits to respond to changes in selective regime are often unknown. Moreover, because most genes expressed in reproductive organs (e.g. testes) are also expressed elsewhere in the body^5,6^, genetic changes that result in reproductive trait modification can potentially lead to negative pleiotropic consequences in either the opposite sex or in other tissues. Despite these constraints, reproductive phenotypes show striking and often rapid divergence, and can promote speciation^7^.

Two closely related *Peromyscus* rodents with highly divergent mating systems show marked variation in male reproductive traits^8−11^. Within the genus, the deer mouse, *P. maniculatus*, is considered one of the most promiscuous species: both sexes mate with multiple partners, often in overlapping series just minutes apart^12^, and females frequently carry multiple-paternity litters in the wild^13^. By contrast, its sister species, the old-field mouse, *P. polionotus*, is strictly monogamous as established from both behavioural^14^ and genetic data^15^. Moreover, relative testes size is roughly three times larger in *P. maniculatus* than in *P. polionotus*^11^, consistent with the well documented relationship between relative testis size and level of sperm competition in rodents^16^. Therefore the competitive environments experienced by sperm of *P. maniculatus* and *P. polionotus* males represent divergent selective regimes.

The factors that regulate mammalian reproductive success are numerous and complex, yet when sperm from multiple males compete for a limited number of ova, the quality of each male’s sperm can influence who will sire offspring^3^. Under intense competition, sperm motility can be a critical determinant of success^17^. Previous studies have shown that *P. maniculatus* sperm swim with greater velocity than *P. polionotus*^9^. A primary energy source for motility is acquired by oxidative phosphorylation in the mitochondria, which are located within the sperm midpiece^18^. The size of the midpiece is thus predicted to positively influence flagellar thrust and sperm velocity^19^ and, indeed, evidence across multiple taxa support the relationship between midpiece size and speed^20−22^.

In this study, we examine the relationship between sperm midpiece length, swimming performance, and reproductive success in *P. maniculatus*, *P. polionotus*, and a hybrid population. We then identify a single gene of large effect that regulates the phenotypic difference in sperm midpiece length between the two focal species, and show how allelic variation at this locus influences sperm swimming velocity and ultimately, male reproductive success.

## Results

### Sperm morphology and performance

We first measured four sperm traits of mice taken from our laboratory colonies of the two focal species, *P. maniculatus* and *P. polionotus* (Fig. 1a). We found that sperm head size does not differ between these species (Fig. 1b-c), but *P. maniculatus* sperm have longer flagella than *P. polionotus* (Fig. 1d; t-test: *P*=8×10^−11^, *df*=9, *n*=10 sperm/male). More specifically, the midpiece region of the flagellum is significantly longer in *P. maniculatus* sperm than in *P. polionotus* (Fig. 1e; t-test: *P*=3×10^−7^*, df*=9, *n*=10 sperm/male). These data are consistent with morphological differences in sperm from wild-caught *P. maniculatus* and *P. polionotus*^10^.

**Figure 1.**
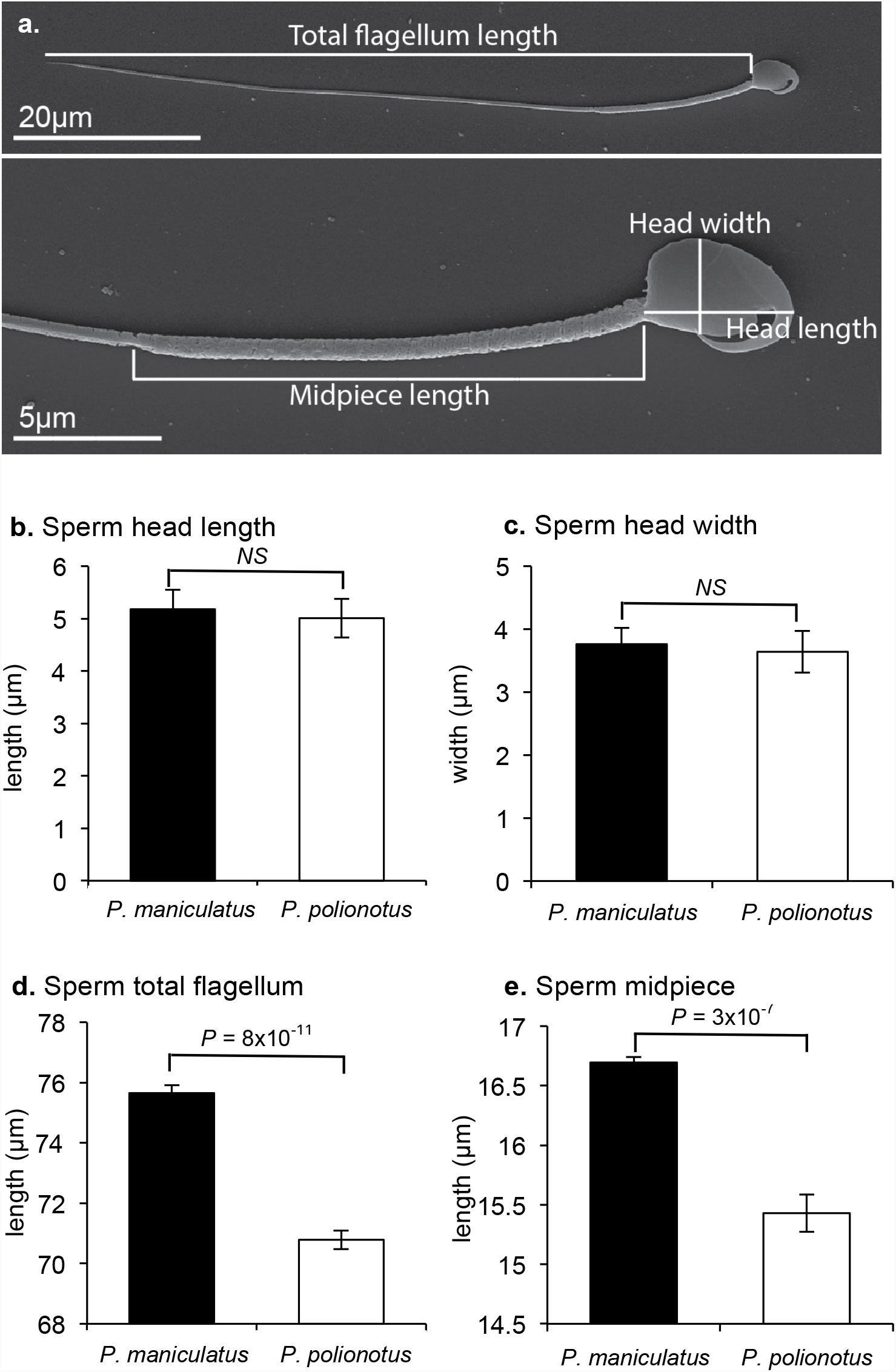
*Peromyscus* sperm morphology. (a) Scanning electron micrographs of a mature *P. maniculatus* sperm cell with morphological features labelled. Mean±SE of *P. maniculatus* and *P. polionotus* sperm (b) head length, (c) head width, (d) total flagellum length and (e) midpiece length (*n=*10 males; *n=*10 sperm/male; t-test). Sperm head and width do not differ significantly, yet *P. maniculatus* total flagellum and midpiece length are significantly longer than those in *P. polionotus* sperm. Note truncated y-axis. *NS* = p-value > 0.05.

Indeed, sperm from the promiscuous *P. maniculatus* males swim with greater velocity (straight-line velocity [VSL]) than sperm of the monogamous *P. polionotus* (t-test: *P*=0.017, *df*=8, *n*=76-549 sperm/male), consistent with our previous results^9^. Two other means of measuring sperm swimming performance, curvilinear velocity (VCL; t-test: *P*=0.0024, *df*=8, *n*=76-549 sperm/male) and average path velocity (AVP; t-test: *P*=0.0039, *df*=8, *n*=76-549 sperm/male) showed a similar difference as in the VSL results. Since all three velocity measures were consistent and are non-independent measurements, we focused on the most conservative estimate, VSL, in subsequent analyses (Supplementary Fig. 1).

To assay the relationship between *Peromyscus* midpiece length and swimming performance in a competitive context, we next conducted a series of swim-up assays, a clinical technique used to screen for highly motile spermatozoa that are most likely to achieve fertilization^23^. We tested sperm with variable midpiece lengths by centrifuging cells and collecting sperm best able to swim towards the surface through a viscous media. We first competed sperm from two heterospecific males (*P. maniculatus* vs. *P. polionotus*), which as a single mixed sample offers the greatest range of sperm morphology, and found that the most motile sperm had a significantly longer midpiece (Fig. 2; t-test: *P*=3.28×10^−4^, *df*=11; *n*=20 sperm/trial). We found a strikingly similar result when the competitions involved sperm from two unrelated conspecific males (*P. maniculatus* vs. *P. maniculatus*), representing a more biologically relevant competition (Fig. 2; t-test: *P*=0.001, *df*=14; *n*=20 sperm/trial), and even within-male competitions (*P. maniculatus*) in which all sperm were harvested from a single male for each trial but still showed variation in midpiece length (Fig. 2; t-test: *P*=0.014, *df*=18; *n*=20 sperm/trial). In total, these results suggest that sperm with larger midpiece regions are more motile, and thus more likely to achieve fertilization^16, 23^, whether they are competing against heterspecific, conspecific or even of other sperm produced by the same male.

**Figure 2.**
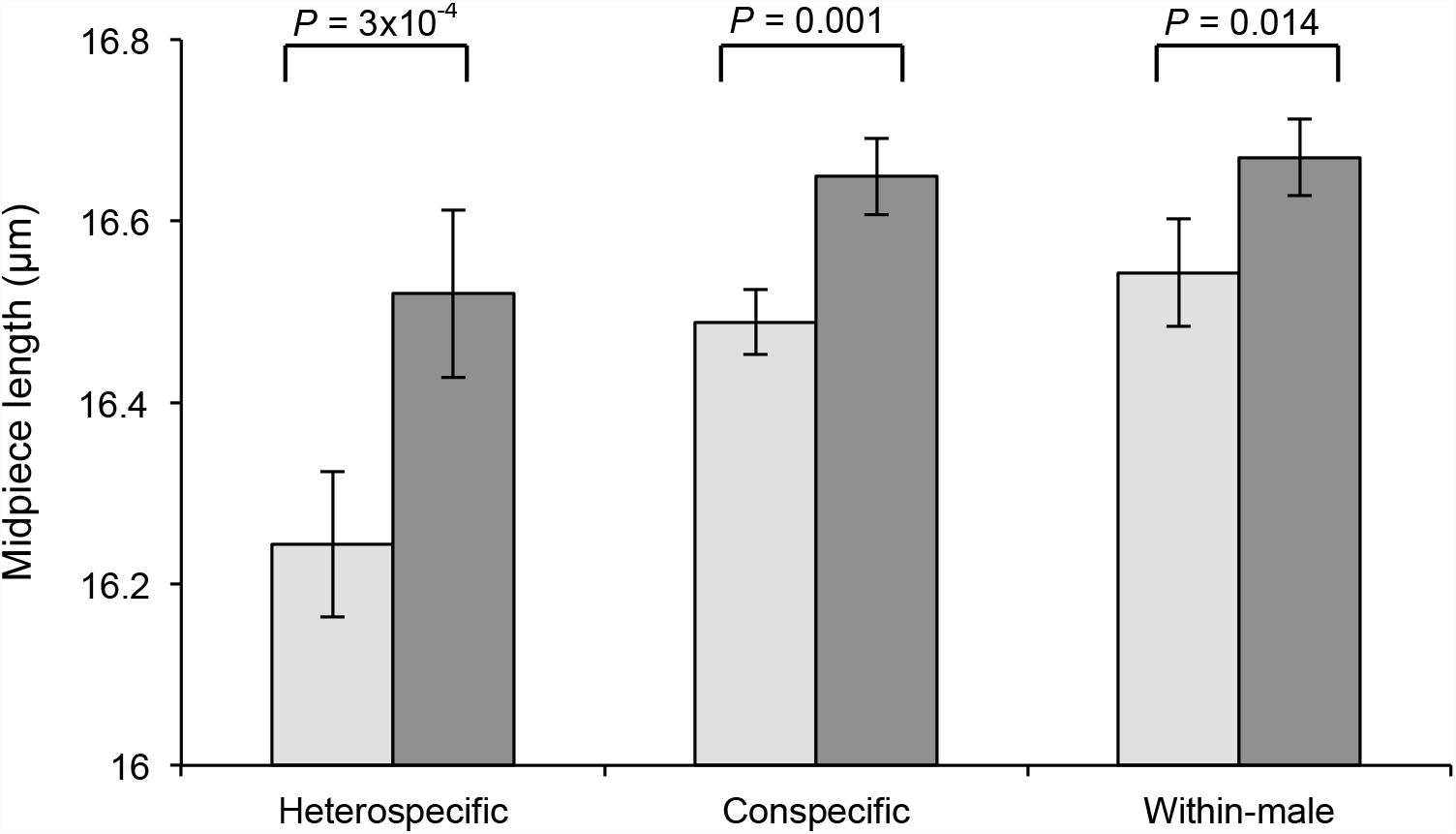
Competitive sperm swim-up assays. Mean±SE of sperm collected prior to spinning (pre-spin; light grey bars) and from the centrifuged sample surface (post-spin; dark grey bars) in heterospecific (*P. maniculatus* vs. *P. polionotus*; *n*=12 trials), conspecific (*P. maniculatus* vs. *P. maniculatus*; *n*=15 trials), and within-male (*P. maniculatus*; *n*=19 trials) competitive swim-up assays. In all three assays, the mean midpiece length (*n=*20 sperm/sample) of competitive sperm collected following centrifugation was significantly longer than those entering (t-test). Note truncated y-axis.

### Genetic mapping of sperm midpiece length

Next, to dissect the genetic basis of adaptive differences in sperm morphology, we performed a genetic intercross between *P. maniculatus* and *P. polionotus* to produce 300 second-generation hybrid (F_2_) male offspring. We then genotyped each F_2_ male at 504 anonymous loci throughout the genome. We identified a single chromosomal region significantly associated with midpiece length variation on linkage group 4 (LG4; Fig. 3; on the basis of logarithm of odds [LOD], significance determined by a genome-wide permutation test with α=0.05). This single region of the genome explains 33% of sperm midpiece length variation in the F_2_ hybrids, and largely recapitulates differences in midpiece length observed between the pure species (Fig. 4). Furthermore, we found that F_2_ males carrying at least one *P. maniculatus* allele at this locus have a significantly longer midpiece than those with none (Fig. 4; t-test: *P*=4.44×10^−15^, *df*=49), suggesting the *P. maniculatus* allele acts in a dominant fashion. Thus, a single large-effect locus explains much of the difference in sperm midpiece length between these two species.

**Figure 3.**
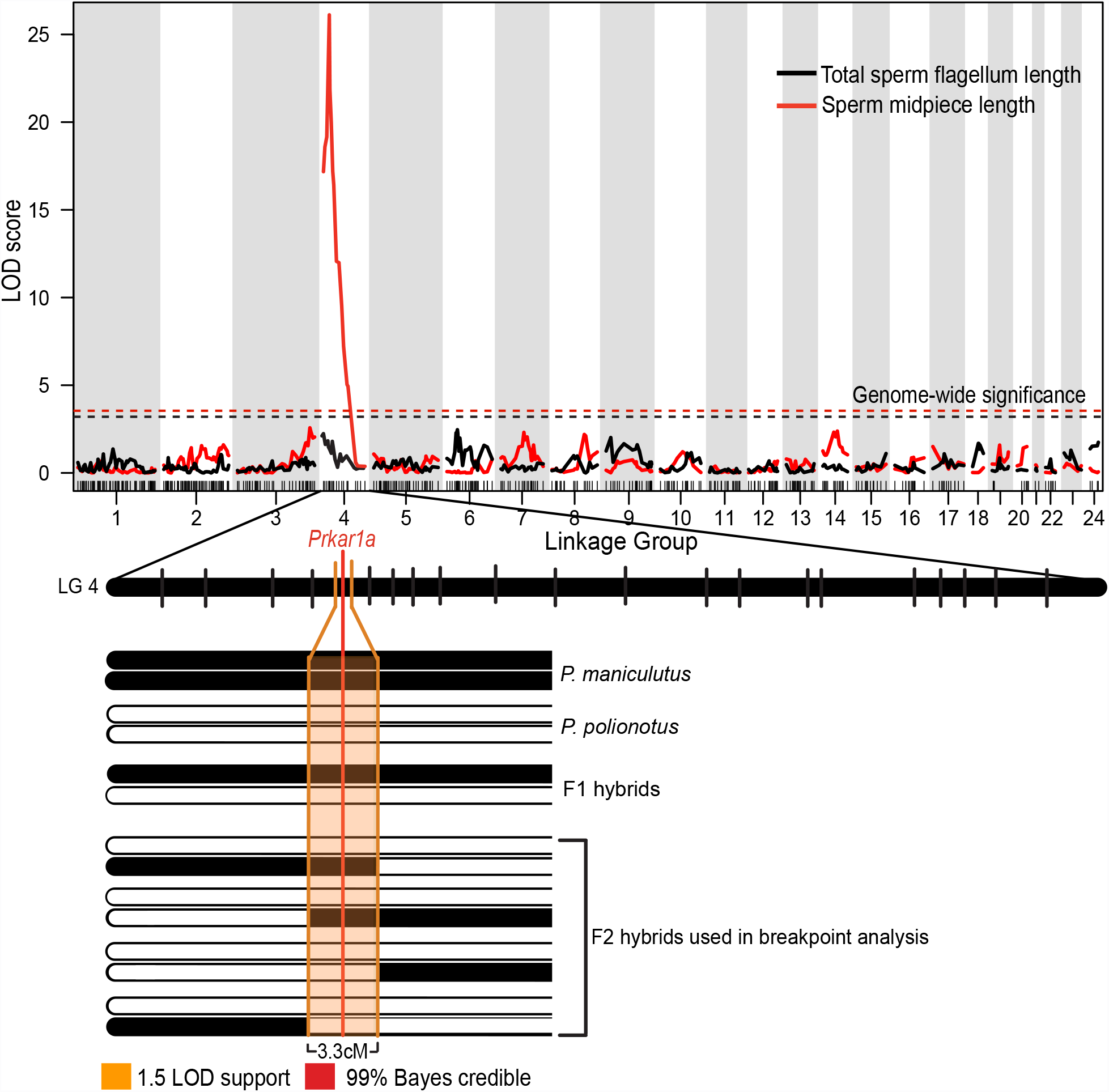
Genetic mapping of sperm morphology. Association between genome-wide SNP markers and total sperm flagellum length (black) and midpiece length (red). Genome-wide significant LOD thresholds for each trait (from 10,000 permutations; α=0.01) indicated by dashed lines. A zoomed in view of Linkage Group 4 (LG4) shows confidence thresholds for QTL associated with midpiece length; 1.5 LOD support interval is shown in orange, the 99% Bayes credible interval is shown; position of the most highly associated gene. *Prkar1a*, is given. F_2_ hybrids with recombination events within the 1.5-LOD support threshold (*n*=11) were used for breakpoint analysis.

**Figure 4.**
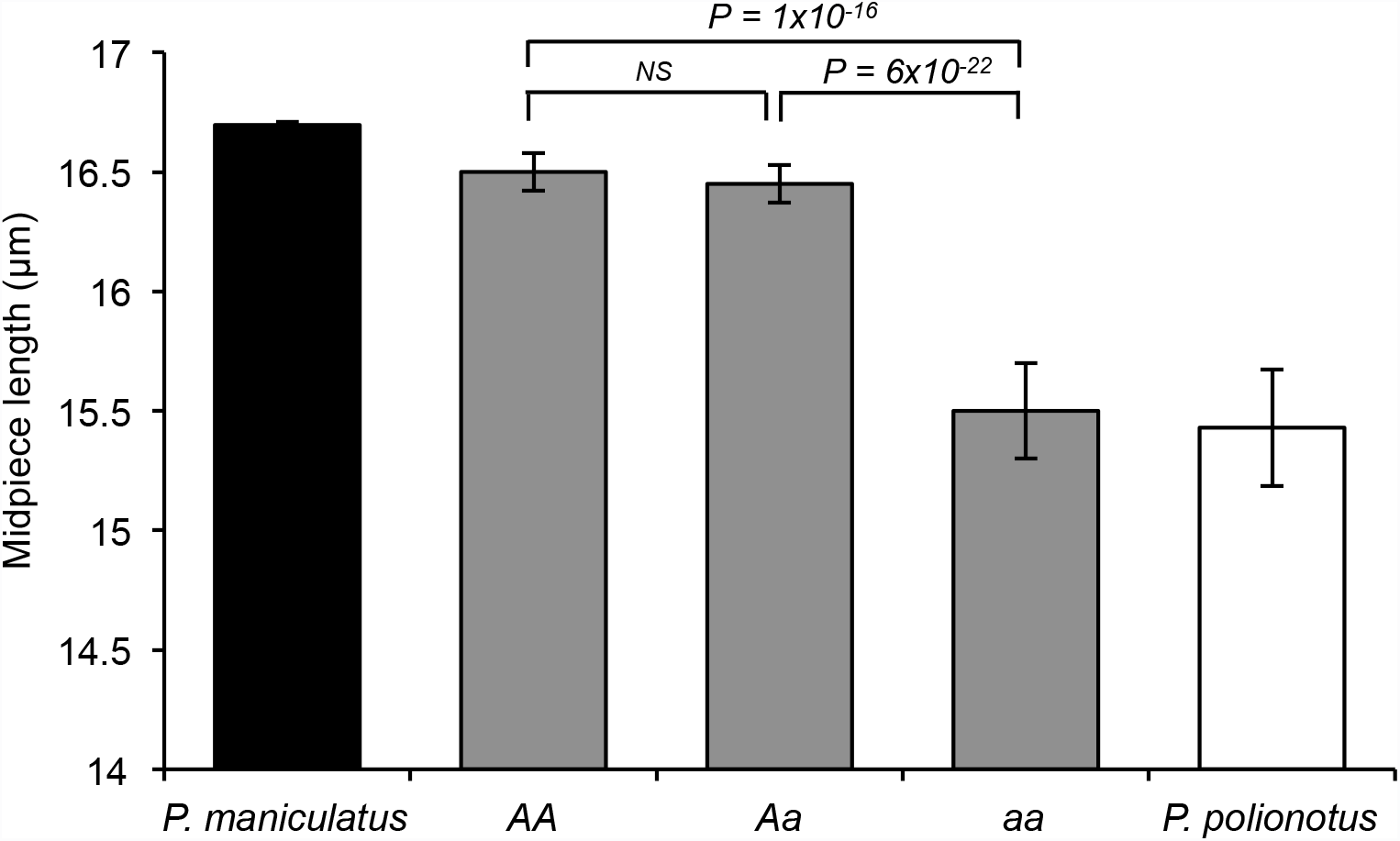
Variation in sperm midpiece length between *Prkar1a* genotypes in hybrids relative to parental phenotypes. Mean±SE midpiece length (*n=*10 sperm/male) of sperm harvested from F_2_ hybrid offspring (*P. maniculatus* [*n*=10; indicated as black bar] x *P. polionotus* [*n*=10; white bar], redrawn from Fig. 1e for reference here). “*AA*” denotes males homozygous for the *P. maniculatus* allele at the *Prkr1a* locus (*n*=61), “*Aa*” for heterozygous males (*n*=130), and “*aa*” for males homozygous for the *P. polionotus* allele (*n=*50). Note truncated y-axis. *NS* = p-value > 0.05, t-test.

While we found a single significant peak associated with sperm midpiece length on linkage group 4 (Fig. 3), we found no significant quantitative trait loci (QTL) for sperm total flagellum length (Fig. 3) or any measure of sperm velocity (VSL, VCL, VAP). The lack of significant QTL for total flagellum length or velocity in this cross does not suggest that variation in these traits lack a genetic basis, rather the result may be due to measurement error or an inability to detect genes of relatively small phenotypic effect, and due to the complex nature of these traits, it is likely that they are controlled by multiple genes. Moreover, velocity is a composite trait likely influenced by sperm morphology as well as other factors. We performed *post-hoc* scans for each of these phenotypic traits (midpiece, total flagellum, VSL, VCL and VAP with 1,000 permutations, α=0.05) with each other trait, and with the genetic marker of highest linkage, as covariates; we found no additional significant QTL.

To further refine the single QTL for sperm midpiece and identify a causal gene, we enriched marker coverage in the 20cM region surrounding the marker of highest association by genotyping each F_2_ male for eight additional single nucleotide polymorphisms (SNPs; Supplementary Table 1). The increased marker density improved our QTL signal and reduced the 1.5-LOD support interval to 3.3cM and the 99% Bayes credible interval to a single locus containing the *Prkar1a* gene (Fig. 3). We then confirmed this association with a genetic breakpoint analysis that included two additional SNPs flanking the marker of highest association, thereby narrowing the 3.3cM interval to 0.8cM around *Prkar1a* (Fig. 5). The *Prkar1a* gene encodes the R1α regulatory subunit of the Protein Kinase A (PKA) holoenzyme and is the only gene within the broader 3.3cM confidence interval previously implicated in male fertility, sperm morphology or sperm motility^24,25^ (Supplementary Table 2). Therefore, *Prkar1a* represents a strong candidate for further functional analyses.

**Figure 5.**
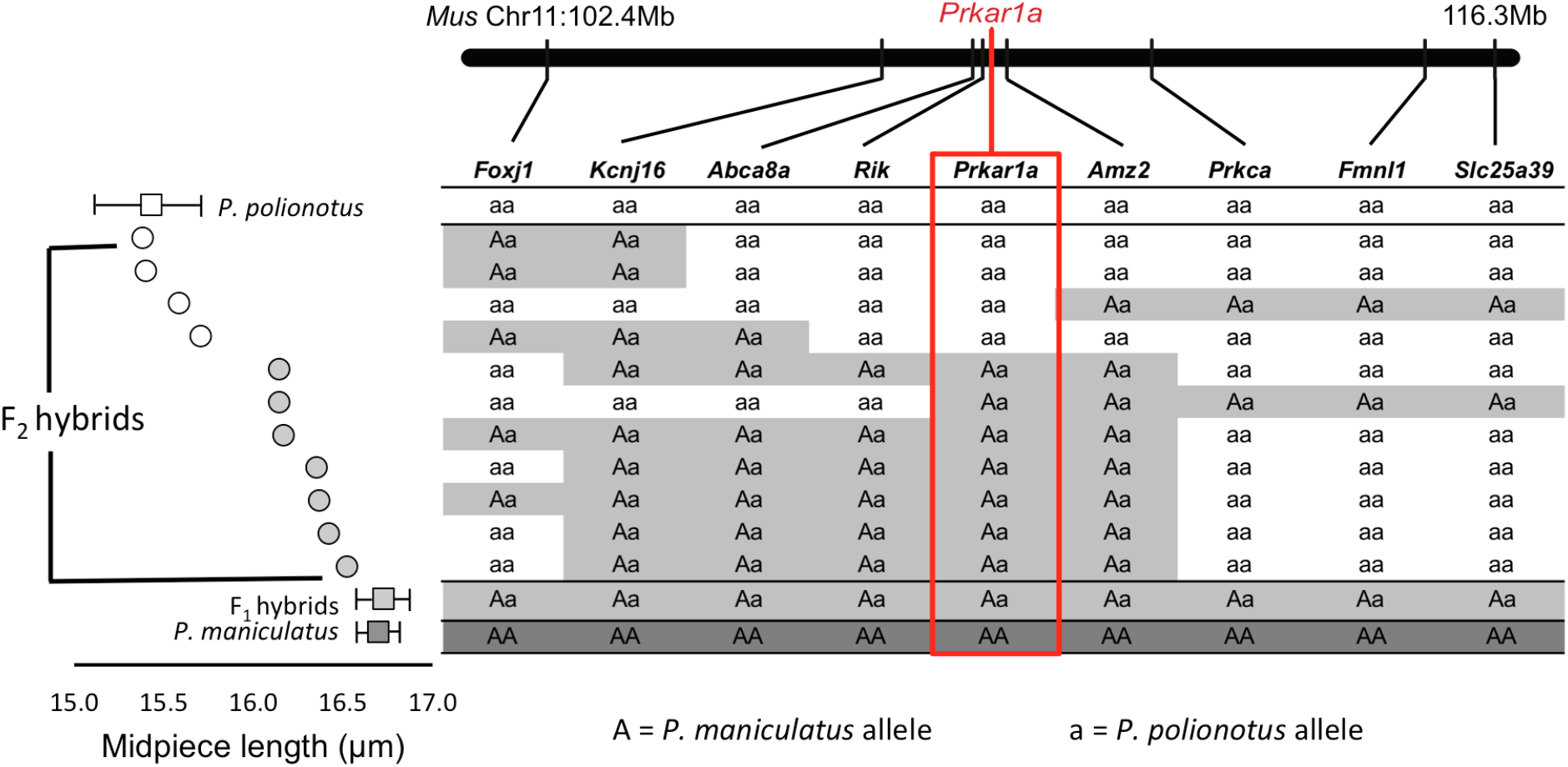
Fine-scale mapping of sperm midpiece length. F_2_ hybrid males (*n*=11) that showed a recombination event in the 1.5 LOD support interval associated with the midpiece QTL (see Fig. 3); males are sorted by mean midpiece length (μm) shown by grey circles (*n*=10 males, *n*=10 sperm/male) from shortest (top) to longest (bottom). Mean±SE midpiece length for pure *P. polionotus, P. maniculatus* (redrawn from Fig. 1) and F_1_ hybrids are shown for reference. Genotypes for each individual at nine markers: “*Aa*” denotes heterozygous genotypes for the *P. maniculatus* and *P. polionotus* alleles, “*aa*” for homozygous *P. polionotus* alleles. Red box highlights the *Prkar1a* genotype that shows a perfect association with midpiece length variation unlike neighboring loci.

### The role of *Prkar1a* in *Peromyscus* reproduction

We found that R1α is abundant and localized in the sperm midpiece in *Peromyscus* (Fig. 6a, Supplementary Fig. 2), consistent with studies in humans^26^ and rats^27^. To confirm the influence of *Prkar1a* on midpiece length, we examined *Mus musculus* C57BL/6 animals with only a single functional copy of the gene (homozygous knockouts are inviable)^28^ and found that the midpiece of *Prkar1a*^+/−^ males is significantly shorter than wild type brothers (Fig. 6b; t-test: *P*=4.54×10^−8^, *df*=7, *n*=20 sperm/male). These findings therefore strongly implicate the *Prkar1a* gene as a major determinant of sperm midpiece length differences observed in *Peromyscus*.

**Figure 6.**
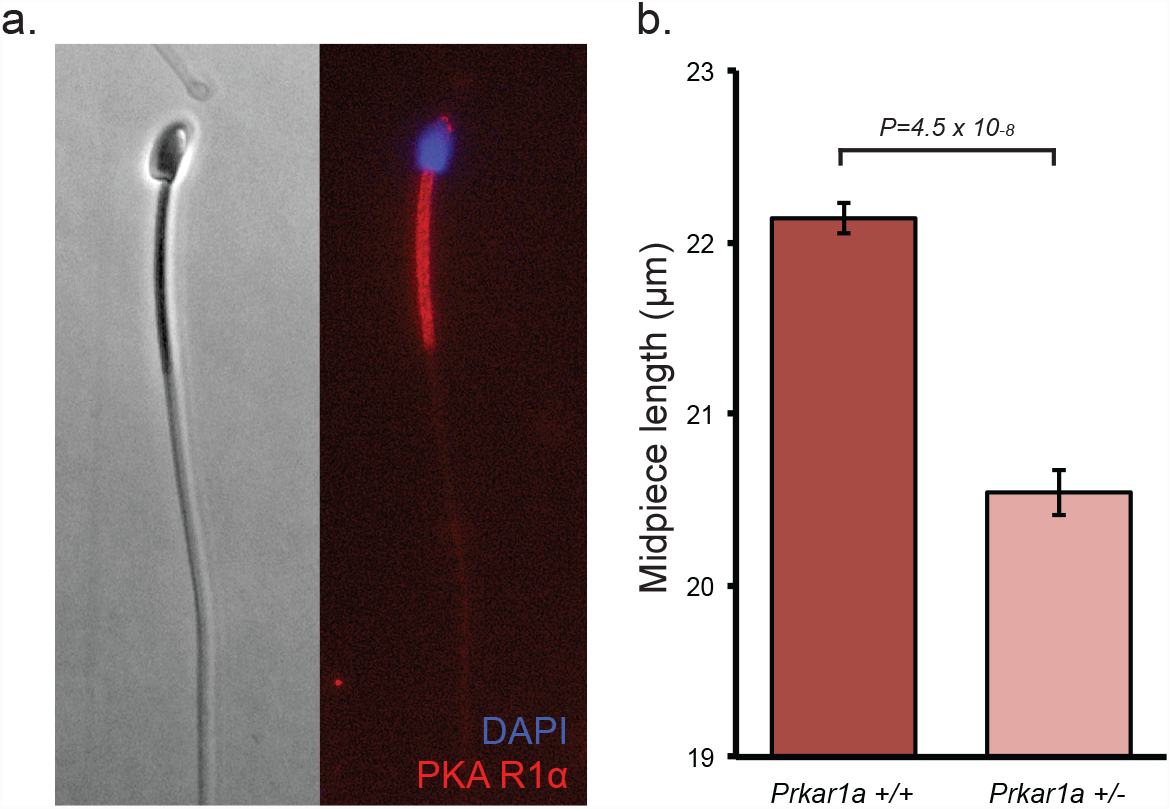
PKA R1α localization and effect on midpiece length. (a) PKA R1α localization in mature *P. maniculatus* sperm cell (400X); immunofluorescence of anti-PKA R1α (red) and DAPI (blue). (b) Mean±SE midpiece length of wild-type *Prkar1a*^+/+^ (red) and heterozygous *Prkar1a*^+/−^ (pink) *Mus musculus* C57BL/6 sperm (t-test: *n*=8 males*, n*=20 sperm/male). Note truncated y-axis.

We next investigated how the PKA R1α subunit differed between our two focal species. First, we found no non-synonymous differences between *P. maniculatus* and *P. polionotus* in the coding exons of *Prkar1a* mRNA (1,146bp), suggesting that both species produce similar R1α proteins. This protein is expressed throughout male germ cell development^29^. In whole testis samples, we found no significant differential expression of *Prkar1a* mRNA and R1α protein levels (mRNA: Supplementary Fig. 3, Bayes factor = 0.071, *Posterior Probability=* 0.008, *n*=8 males; protein: Fig. 7; t-test: *P*=0.664, *n*=5 males), possibly because testis samples contain cells of multiple developmental stages. However in mature, fully developed sperm released from the caudal epididymis, we found *P. polionotus* express significantly more R1α protein than *P. maniculatus* (Fig. 7; t-test: *P*=0.00235, *n*=6 males). These data, in combination with the knockout phenotype in *Mus*, suggest that changes in the expression of R1α can lead to changes in sperm midpiece length.

**Figure 7.**
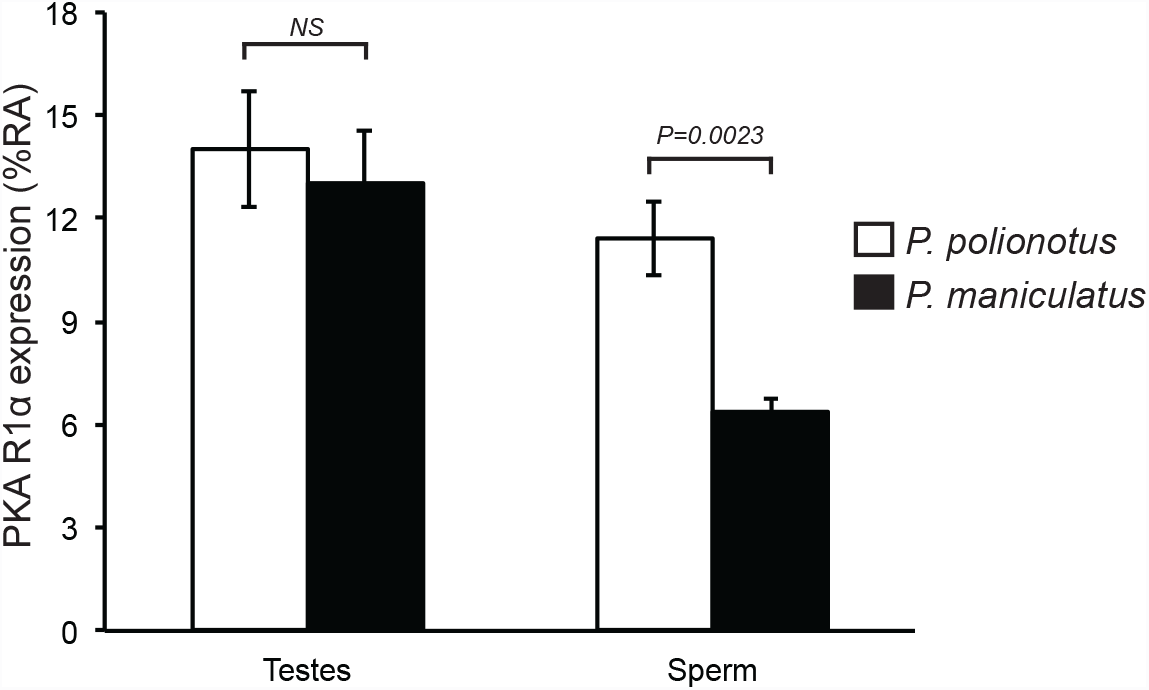
PKA R1α expression. (c) PKA R1α protein expression as measured by percentage of relative abundance (%RA) in testes (*n*=5) and epididymal sperm (*n*=6) samples from *P. polionotus* (white bars) and *P. maniculatus* (black bars) males. *NS* = p-value > 0.05, t-test.

### Linking genotype, phenotype and fitness in a hybrid population

In addition to genetic mapping, our genetically heterogeneous F_2_ hybrid population also enables us to test for statistical associations among traits. First, to test the prediction that sperm midpiece length influences swimming performance, we compared the mean straightline velocity (VSL) to both mean midpiece length and flagellum lengths of sperm from F_2_ males using a linear regression, with each trait as the independent variable. While we found no significant association with flagellum length [*R^2^*=0.003, *P*=0.29, *df*=232], midpiece and VSL show a significant positive correlation [Fig. 8a; *R^2^*=0.028, *P*=0.0057, *df*=232], which remains significant after considering flagellum length as a covariate [*R^2^*=0.028, *P*=0.012, *df*=232, midpiece was the only coefficient with *P*<0.05]). Moreover, we found no significant association between midpiece length and total flagellum length in F_2_ males (*R^2^*=0.004, *P*=0.14, *df*=232), which suggests that these two traits are genetically independent. These results show that VSL is correlated with midpiece length in *Peromyscus*, either because increase in midpiece length leads to increased speed (i.e., variation in these two traits share a pleiotropic genetic basis), or less likely, these two traits are influenced independently by tightly linked genes.

**Figure 8.**
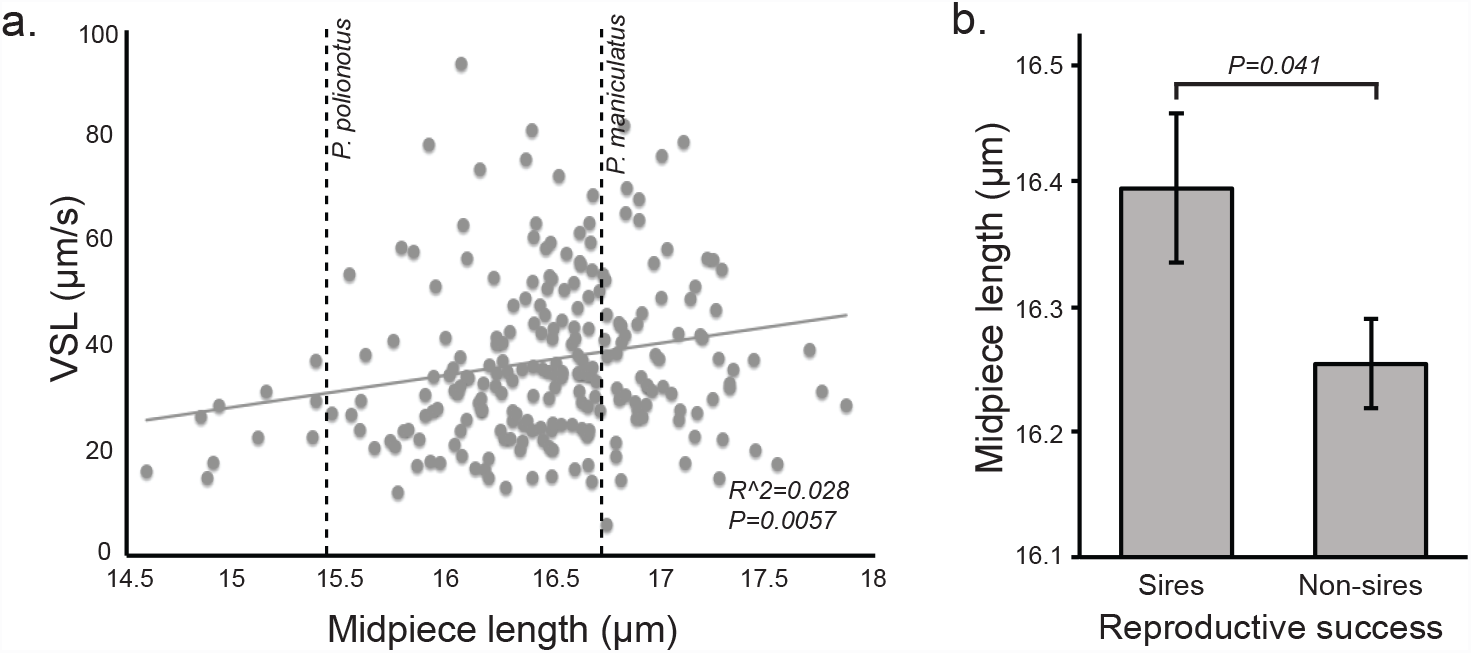
Sperm performance and male fertility in F_2_ hybrids. (a) Association between midpiece length and straight-line velocity of F_2_ hybrid males (linear regression: *n*=233 males, *n*=10 sperm/male). For reference, midpiece length of each parental species is plotted as a dashed line. (b) Mean±SE sperm midpiece length of F_2_ males that did (*n*=85) and did not (*n*=173) sire offspring (t-test). Note truncated y-axis.

We next examined how variation at the *Prkar1a* locus predicts sperm morphology and performance in the F_2_ hybrid population. We found that F_2_ males carrying at least one *P. maniculatus Prkar1a* allele (*AA* or *Aa*, as defined by the *Prkar1a* SNP) produce sperm with significantly longer midpiece, as mentioned earlier (Fig. 4), but their sperm also swim with greater velocity (VSL) than sperm from males homozygous for *P. polionotus Prkar1a* allele (*aa*; mean VSL±SE: *AA*=78.7±1.9µm/s, *Aa*=77.5±1.1µm/s, *aa*=73.1±1.8µm/s, t-test: *P*_*AA*-*aa*_=0.040, *P*_*Aa*-*aa*_=0.044, *df*=49). Therefore, *Prkar1a* genotype is associated with both sperm morphology and swimming performance.

Finally, to understand how allelic and phenotypic variation influence male reproductive success, we scored which F_2_ males sired offspring in natural matings. We found that F_2_ hybrid males carrying at least one dominant *P. maniculatus Prkar1a* allele were more likely to sire offspring when paired with a female for two weeks than those homozygous for the *P. polionotus* allele (χ^*2*^_AA-Aa-aa_=6.35, *P*=0.042, *df*=2). This is a conservative estimate of male fertility because many false negatives were likely included due to pairs that failed to mate, infanticide, female infertility and other unrelated physiological or behavioural conditions of the animals, which make an effect more difficult to detect. Moreover, this result is consistent with lower reproductive success in pure *P. polionotus* matings compared to *P. maniculatus* under similar conditions (*z*=1.37, *P*=0.0071, *df*=19). Furthermore, F_2_ males who sired pups had sperm with significantly longer midpiece regions than those that did not reproduce (Fig. 8b; t-test: *P*=0.041, *df*=84). Together, these analyses show that males carrying at least one copy of the *P. maniculatus Prkar1a* allele produce significantly faster sperm with longer midpieces, and also benefit from greater reproductive success, suggesting a direct link between fitness, phenotype and genetic variation at the *Prkar1a* locus.

## Discussion

When females mate with multiple partners within a reproductive cycle, males can continue to compete for reproductive success long after mating has occurred as their sperm compete for the fertilization of a limited number of ova. The strength of post-copulatory sexual selection is therefore largely determined by female mating strategy. In this study, we examined two species of *Peromyscus* mice with divergent mating systems. Female *P. maniculatus* mate with multiple males in a reproductive cycle, allowing for sperm of different males to compete within the female reproductive tract for fertilization success; in contrast, *P. polionotus* females mate monogamously and thus sperm competition is limited. Theory predicts that these differing competitive regimes may favour the evolution of trait divergence. Our results suggest that the difference observed in sperm midpiece length between *P. maniculatus* and *P. polionotus*, confers an important reproductive advantage that improves sperm swimming performance. We first found that sperm with longer midpiece are more motile in a competitive swim-up assay. Second, within our hybrid population, we observed a positive relationship among sperm midpiece length, swimming velocity and male reproductive success. The targets of post-copulatory sexual selection can vary tremendously across taxa, but in this system, our results are consistent with the hypothesis that selection favors sperm with longer midpiece regions, consistent with findings in other species^20−22^.

If a simple relationship between midpiece length and fitness exists, it is puzzling why a monogamous species would not share the same sperm morphology as its promiscuous sister-species. The functional relevance of the midpiece, in both evolutionary and human fertility studies, is controversial^30,31^, and many closely-related species vary extensively in this trait^32−34^. Nonetheless, while drift and/or selection acting on pleiotropic traits could lead to shorter midpiece, sperm cells with more or larger mitochondria afforded by the larger midpiece also may experience greater oxidative stress, which is known to increase mutagenesis in germ cells^35^. Therefore, when sperm competition is absent, such as in *P. polionotus*, the benefits conferred by producing faster sperm may not outweigh the associated costs. However in the highly competitive environment that *P. maniculatus* sperm experience, even small increases in sperm performance could differentiate those that reproduce and those that do not^7^. Thus, the balance between negative and positive effects of cellular respiration in sperm may determine the relative roles of natural and sexual selection as drivers of interspecific midpiece variation seen in *Peromyscus*, and across animals more generally.

Using a forward-genetics approach, we identified a single gene of large effect on midpiece size, *Prkar1a*, which encodes for the R1α regulatory subunit of the Protein Kinase A. We then corroborated the role of this gene by demonstrating that lab mice with a dominant negative mutation in *Prkar1a* have shorter midpiece than their wild-type brothers. Successful fertilization requires precise temporal and spatial regulation of PKA activity, the main downstream effector of cellular cyclic AMP concentrations, and is in part, regulated by R1α^29^. The R1α subunit is known to influence gross sperm morphology, motility and fertility in humans presenting Carney Complex (a disease associated with mutations in *Prkar1a*) and in *Mus musculus Prkar1a* mutants^24^. While most reports on *Prkar1a* implicate negative consequences for male fertility and are associated with large-scale changes in R1α expression^24,29,36–38^, our results suggest that the subtle tuning of the expression of this regulatory subunit may also confer beneficial effects. In *Peromyscus*, we found that R1α is similarly abundant in the testes of the two focal species, yet is differentially expressed in mature sperm of *P. maniculatus* and *P. polionotus*. Considering the heterogeneous nature of testis tissue contains germ cells at various stages of their differentiation process and Sertoli cells, it is likely that subtle difference in *Prkar1a* mRNA abundance or R1α protein may not be detectable if expressed in a limited number of cell types. Our analysis of mature spermatozoa, however, suggests that cell type-specific R1α expression differences present during later stages of *Peromyscus* spermatogenesis are likely to regulate midpiece length.

The rapid evolution of reproductive protein coding regions is a well-known response to post-copulatory sexual selection^39^; we demonstrate here that expression changes in gametes of a broadly-expressed gene can also be a target of selection. Over 50% of mammalian genes are expressed in the testis and most of these genes are also expressed in other tissues^5,6^; therefore, cell-type specific changes in protein expression in reproductive tissues are likely to be a common mechanism by which selection in males can operate with swiftness and without deleterious effects in females or other tissues.

## Methods

### Mice

Wild derived *Peromyscus maniculatus bairdii* and *Peromyscus polionotus subgriseus* were originally obtained from the Peromyscus Genetic Stock Center at the University of South Carolina and have been maintained at Harvard University in accordance with guidelines established by Harvard’s Institutional Animal Care and Use Committee. Adult sexually-mature *P. polionotus* and *P. maniculatus* males were used to collect data for cross-species comparisons. In addition, we bred four mice, two *P. polionotus* males and two *P. maniculatus* females, to produce 40 first-generation (F_1_) hybrids, and then intercrossed siblings to generate second-generation (F_2_) hybrid progeny. For genetic mapping, we used 300 F_2_ hybrid males and obtained genotypic and phenotypic data as described below. All males were sexually mature and were paired with a female prior to harvesting sperm.

*Mus musculus* females heterozygous for the *Prkar1a* locus on a C57BL/6 background^28^ were mated to wild type C57BL/6 males to produce wild type and *Prkar1a*^+/−^ offspring. These mice were a generous gift of Dr. Stanley McKnight at the University of Washington.

### Sperm Analysis

After sacrifice via carbon dioxide overdose, we immediately removed the left caudal epididymis of each F_2_ male with a single cut at the intersection of the epididymis and the vas deferens. We then submersed the tissue in 1 mL of warmed Biggers-Whitten-Whittingham (BWW) medium^40^, and incubated the tissue for 10 min at 37°C to release motile sperm. We then removed the epidydimal tissue, gently swirled the solution, placed 20µl of medium containing live sperm on a plastic microscope slide and covered the sample with a plastic coverslip (plastic reduces adhesion of sperm to the slide compared with glass products). We recorded 5 sec videos of live sperm at 100X magnification under phase contrast conditions on an upright microscope (AxioImager.A1, Zeiss, Jena, Germany). We then acquired sperm swimming performance data from the videos using the Computer Assisted Sperm Analyzer plugin for NIH ImageJ^41^ by measuring each motile cell in the frame (*n*=4-549 cells/male) to calculate the mean straight-line velocity (VSL), curvilinear velocity (VCL) and average path velocity (VAP). Cells aggregated into groups, dead sperm cells and those with a beating flagellum but stuck to the slide were excluded from the analysis. Sperm collected from *P. maniculatus* achieved higher velocity than *P. polionotus* sperm on all three measures (Supplementary Fig. 1; t-test: *P*_*VSL*_=0.017, *P*_*VCL*_=0.0024, *P*_*VAP*_=0.0039, *n*_*male*_=9).

We preserved the remaining sperm cells in 2% paraformaldehyde (Sigma-Aldrich, St. Louis, MO) for morphological analysis. Following fixation, we spread 20µl of suspended sperm on a glass microscope slide and mounted the cells in Fluoromount-G (Southern Biotech, Birmingham, AL). We imaged 10 sperm showing no obvious abnormalities from each male at 400X magnification under phase contrast conditions on an upright microscope (AxioImager.A1), consistent with standard practice^34,42–44^. For each sperm cell from *P. maniculatus* and *P. polionotus* males we measured (1) sperm head length (the longest region of the sperm head), (2) sperm head width (the widest region of the head perpendicular to the length measure), (3) total sperm flagellum length (including midpiece, principal piece and terminal piece of cell), and (4) midpiece length (from the base of the sperm head to the midpiece-principal piece boundary; see Fig. 1a) using the curve-spline tool in the AxioVision Image Analysis Software (Zeiss, Jena, Germany). Intraclass correlation coefficients among sperm within F_2_ hybrid males were significant and high (midpiece=0.651, *P*<0.05; flagellum=0.612, *P*<0.05), suggesting repeatability of the measure and substantial heritability of sperm morphology, as expected from the significant differences in parental species means. *Mus musculus* C57BL/6 wild type and *Prkar1a*^+/−^ males’ sperm midpiece regions were measured and analyzed identically to *Peromyscus* sperm.

We compared the sample means of pure *P. maniculatus* and *P. polionotus* sperm morphology (head length, head width, total flagellum length, midpiece length) and velocity (VSL, VCL, VAP) using two-tailed unpaired t-tests and adjusted alpha for multiple comparisons using Bonferroni correction.

F_2_ hybrid males that did not produce any sperm were excluded from all analyses incorporating sperm velocity or morphology measures, including genetic mapping; however, we used the genotypic information from these males in linkage map construction (see below). All analyses were performed in R statistical software^45^.

### Fitness Assays

We tested for an effect of sperm midpiece length on competitive success by performing three types of sperm swim-up assay: heterospecific competitions including the sperm of a *P. maniculatus* and a *P. polionotus* male, conspecific competitions between the sperm of two *P. maniculatus* males, and within-male competitions between the sperm of a *P. maniculatus*, including replicates of each assay. We collected live sperm (following methods above) and placed approximately equal concentrations from the two donors (for within-male assays, we added twice the volume of a similar concentration) in 0.5mL 10% polyvinylpyrrolidone in BWW medium. We took a sample of the sperm mixture (“pre-spin”) and centrifuged the mixture at 150g for 10min at 37°C to pellet cells. We then collected a sample of cells just below the surface of the medium following centrifugation (“post-spin”) and measured midpiece length of sperm in each sample (*n*=20sperm/sample). The cells near the top of the tube after spinning are those that are capable of rapidly swimming up through a viscous media. To analyze the results from these assays, we used two-tailed paired t-tests with Bonferroni correction to compare the mean midpiece size of a sample of sperm that entered the competition (the pre-spin sample) to the sperm that were able to swim to the top of the tube after centrifugation (the post-spin sample). This technique is commonly used to screen for the highly motile sperm most likely to achieve fertilization^23,46^.

To test for an effect of sperm morphology on reproductive success we weaned each of the F_2_ hybrid males from their parents at 25 days of age and housed them with same-sex littermates until they were at least 68 days old and sexually mature. We then housed each male with an F_2_ female chosen at random from the same grandparents as the male for at least 7 days (range 7-22 days; we found no effect of pairing length on reproductive success: unpaired two-tailed t-test comparing pairing length of sires and non-sires, *P*=0.34, *df*=84). This time paired with a female reduced male phenotypic variability that might arise from dominance hierarchies among male littermates and to minimized differences in reproductive condition among males by exposing all to a female in oestrus (*Peromyscus* estrous cycle is 5 days^47^). Finally, we sacrificed males at 80-169 days of age to harvest sperm (we found no effect of male age on reproductive success: unpaired two-tailed t-test comparing mean age of sires and non-sires, *P*=0.69, *df*=84), and recorded any observed offspring that resulted from the pairing of these F_2_ males with F_2_ females.

### Genetic Analysis

We extracted genomic DNA from liver tissue using either phenol chloroform (Automated DNA Extraction Kit, AutoGen, Holliston, MA) or DNeasy Kits (Blood & Tissue Kit, Qiagen, Hilden, Germany). We identified single nucleotide polymorphisms (SNPs) and assigned genotypes for each individual by double digest Restriction-site Associated DNA Sequencing (ddRADseq)^48^. Briefly, we digested ~1µg of gDNA for each individual with two restriction endonucleases: EcoR1 and Msp1 (New England BioLabs, Ipswich, MA). We ligated the resulting fragments to sequencing adapters containing a unique barcode for each individual male. We then pooled these barcoded fragments from multiple individuals and isolated the fragments in the size range of 280-320bp using a Pippin Prep (Sage BioSciences, Beverly, MA). Finally, we amplified the remaining fragments using a Phusion High Fidelity PCR Kit (Thermo-Fisher, Waltham, MA) and sequenced the resulting libraries on a Genome Analyzer IIx or a HiSeq 2500 (Illumina, San Diego, CA). We recovered 1753 informative SNP markers that consistently differed between the two parental strains used to generate the hybrid population, and which were confirmed as heterozygous in the first generation hybrids (F_1_). We then pruned our marker set to exclude any markers genotyped in fewer than 100 individuals or with genotype information identical to another marker.

We conducted all genetic mapping analyses using R/qtl software^49^, an add-on package for R statistical software ^45^. To construct a genetic linkage map, we calculated linkage distances based on the fraction of recombination events and Logarithm of Odds (LOD) scores between all SNP marker pairs. Next, we grouped markers by varying the recombination parameters until we recovered a map with 23 linkage groups containing at least 5 markers each (the karyotypes of both species are known [n=24chromosomes], however we are unable to recover the majority of the Y chromosome with the cross design employed in this study). Any markers not included in the 23 linkage groups were excluded. Finally, we refined this map by ordering the markers within each linkage group in overlapping windows of 8 markers and minimizing the frequency of recombination events between markers in each window. The resulting genetic linkage map contained 504 in 23 linkage groups.

To identify quantitative trait loci (QTL) contributing to sperm morphology, we performed Haley-Knott regression and interval mapping analyses sequentially with 1,000 permutations and α=0.05 in R/qtl^49^. We found a found a single genomic region with a significant association for sperm midpiece length on LG4 Fig. 3); LG4 is syntenic to *Mus musculus* chromosome 11 and *Rattus norvegicus* chromosome 10^50^. To narrow the QTL interval, we genotyped the F_2_ males for additional markers within the 20cM region of the genome surrounding the marker most strongly associated with sperm midpiece length (Supplementary Table 1). This region of interest is syntenic to chromosome 11:97–114Mb in *M. musculus* and chromosome 10:91–104Mb in *R. norvegicus*^51^. We designed markers in eight genes – *Foxj1, Kcnj16, Prkar1a, Prkca, Fmnl1, Klhl10, Slc25a39*, and *Ttll6* – based on position and prioritized those implicated in male fertility, spermatogenesis, cytoskeletal organization, mitochondrial function, and/or specifically or highly expressed in the testes or during spermatogenesis^51−53^. We genotyped all F_2_ males for informative SNPs residing in each of these genes using custom designed TaqMan SNP Genotyping Assays (Thermo-Fisher, Waltham, MA), except in the case of *Ttll6*, where we used a restriction enzyme digest assay (all primers and TaqMan probe sequences are available in Supplementary Table 1; methods followed manufacturer’s instructions). The genotype data from these 8 SNP markers helped to better refine the QTL for sperm midpiece length and increased the association between genotype and phenotype (peak LOD score=25.7 [genome-wide LOD threshold=3.69 with 10,000 permutations and α=0.01]). Using two measures of confidence, we found that midpiece length mapped to (1) a single marker residing in the gene, *Prkar1a*, based on the 99% Bayes credible interval, and (2) to a 3.3cM region surrounding *Prkar1a* based on 1.5 LOD support interval^49^. We then PCR-amplified (primers available in Supplementary Table 1) and sequenced ~1 kb of gDNA fragments within the genes *Abca8a* and *Amz2* in the parents of the cross and all F_2_ males that showed a recombination event in the surrounding region (*n*=13; Fig. 5). On the basis of midpiece length of these F_2_ recombinants and their genotypes at the two additional markers, we were further able to narrow the QTL to 0.8cM by excluding association with *Amz2* and *Abca8a* (Fig. 5) and confirm the association between midpiece length and *Prkar1a*.

To investigate the association between reproductive success and genotype, we compared the frequencies of F_2_ males homozygous for the *P. maniculatus Prkar1a* allele (*AA*), the *P. polionotus* allele (*aa*), or heterozygous (*Aa*) who sired offspring using a 3x2 Chi-squared contingency table (χ^*2*^_AA-Aa-aa_=6.35, *P*=0.042, *df*=2).

To investigate genetic differences between *P. maniculatus* and *P. polionotus* within *Prkar1a*, we sequenced the protein-coding region by extracting mRNA from the decapsulated testis tissue of *P. maniculatus* and *P. polionotus* males using an RNeasy Mini Kit (Qiagen, Hilden, Germany), converting the mRNA to cDNA using SuperScript II Reverse Transcriptase (Thermo-Fisher, Waltham, MA) and a poly-T primer (T16). We amplified the entire coding sequence (1,146bp) using two pairs of primers (named by location in *Prkar1a*): Exon1c_F: 5’GCCATGGTTCCTCTGTCTTG3’, Exon8_R: 5’AGAACTCATCCCCTGGCTCT3’; Exon7_F: 5’ATGTGAAACTGTGGGGCATT3’ and 3’UTR_R: 5’ACGACCCAGTACTTGCCATC3’.

To estimate *Prkar1a* mRNA abundance in whole testes of *P. maniculatus* and *P. polionotus (n=8)*, we conducted an RNAseq experiment. Specifically, we extracted total RNA using the TRIzol reagent (Thermo-Fisher, Waltham, MA), purified it with RNeasy Mini Kit (Qiagen, Hilden, Germany), and constructed cDNA libraries using the TruSeq RNA v2 Kit (Illumina, San Diego, CA) following manufacturer instructions. Multiplexed libraries were sequenced on an Illumina HiSeq2500 (Illumina, San Diego, CA) in paired-end mode with a read length of 150bp (average read depth=30.4million reads/sample). We removed low quality and adaptor bases from raw reads using TrimGalore v0.4.0 (http://www.bioinformatics.babraham.ac.uk/projects/trim_galore/) and mapped the trimmed reads to the genome using the STAR RNA-Seq aligner v2.4.2a in two-pass mode^54^. We next obtained the *P. maniculatus* genome sequence and annotation from the Peromyscus Genome Project (Baylor College of Medicine; https://www.hgsc.bcm.edu/peromyscus-genome-project) and created a *P. polionotus* genome and annotation by incorporating SNPs and indels into the *P. maniculatus* Pman_1.0 reference. We estimated expression using STAR alignments in transcriptomic coordinates and the MMSEQ package v1.0.8^55^, and calculated differential expression using mmdiff^56^.

To validate the *Prkar1a* mRNA estimates from RNAseq, we conducted quantitative PCR (qPCR) on testis of *P. maniculatus* and *P. polionotus*. First, we extracted mRNA from whole testes using an RNeasy Mini Kit, converted the mRNA to cDNA with SuperScript II Reverse Transcriptase (Thermo-Fisher, Waltham, MA) using a poly-T primer (T16). We then amplified a fragment with identical primer-binding sites *P. maniculatus* and *P. polionotus* using KAPA SYBR^®^ FAST Universal 2X qPCR Master Mix (Kapa Biosystems, Wilmington, MA) and the following primers: 5’TCCAGAAGCACAACATCCAG3’ and 5’TTCATCCTCCCTGGAGTCAG3’. We conducted assays in triplicate, calculated ∆CT values with *TATA Binding Protein* (*TBP;* 5’CTCCCTTGTACCCTTCACCA3’ and 5’GAAGCGCAATGGTCTTTAGG3’), a standard control gene and validated with RNAseq. To test for significant differences in RNA abundance, between *P. maniculatus* and *P. polionotus* males, we used two-tailed unpaired t-tests.

### Protein Analysis

To localize the expression of PKA R1α, we fixed epididymal sperm in 2% paraformaldehyde and 1.25% glutaraldehyde on a microscope slide for 15min. We then washed the cells in phosphate-buffered saline with 0.1% Tween 20 (PBT) for 15min, and blocked in PBT with 3% bovine serum albumin (BSA) for 1hr at room temperature (RT). Next we incubated the cells overnight at 4°C with the primary antibody, PKA R1α (Santa Cruz Biotechnology #18800, Dallas, TX), which we diluted 1:100 in PBT with 3% BSA. The following day we washed cells in PBT for 1hr at RT 3 times, then incubated with the secondary antibody, Alexa Fluor 546 (Thermo-Fisher #A11056, Waltham, MA), diluted 1:500 in PBT with 3% BSA, at 1:1000 for 1hr at RT. Cells were then washed in PBT for 1hr at RT 3 times, stained with DAPI (Thermo-Fisher, Waltham, MA) to visualize DNA within cells for 15min at RT, washed a final time in PBT for 15min, and mounted in Fluoromount-G (Southern Biotech, Birmingham, AL). In addition, we controlled for non-specific binding of the secondary antibody by performing a side by side comparison with cells processed identically to the above methods except that instead of treating with the primary antibody, cells were solely treated with PBT with 3% BSA, the secondary antibody and DAPI. We viewed cells at 400X and 1000X magnification on an upright microscope (AxioImager.A1, Zeiss, Jena, Germany).

After sacrifice via carbon dioxide overdose, we removed one testis and stored one at −80°C, and submersed both caudal epididymes in 1mL of Modified Sperm Washing Medium (Irvine Scientific, Santa Ana, CA) at 37°C for 10min to release motile sperm. We then removed the epididymes and incubated sperm for an additional 45min at 37°C before pelleting them at 12,000g for 5min. We washed sperm cells in phosphate buffered saline twice, and stored the sample at −80°C. We lysed samples at 50Hz for 2min in lysis buffer (8M Urea, 1% SDS, 50mM Tris pH 8.5, cOmplete Protease Inhibitor [Roche], PhosSTOP [Roche]), rocked for 30 min and centrifuged 12,000g for 20min, all at 4°C. Samples were then processed for multiplexed quantitative mass spectrometry analysis and analyzed through the Thermo Fisher Scientific Center for Multiplexed Proteomics at Harvard Medical School. Samples were subjected to tandem protein digestion using trypsin and Lys-C protease (Thermo-Fisher, Waltham, MA), peptide labeling with Tandem Mass Tag (TMT) 10-plex reagents and peptide fractionation into 12 fractions. Multiplexed quantitative mass spectrometry data were collected on an Orbitrap Fusion mass spectrometer operating in a MS3 mode using synchronous precursor selection for the MS2 to MS3 fragmentation^57^. MS/MS data were searched with SEQUEST against a custom *Peromyscus* database with both the forward and reverse sequences, and included controlling peptide and protein level false discovery rates, assembling proteins from peptides, and protein quantification from peptides. We compared the relative abundance PKA R1α, normalized across all TMT reporter channels, in *P. maniculatus* and *P. polionotus* sperm and testes using two-tailed unpaired t-tests.

## Acknowledgements

We thank S. McKnight for generously donating Prkar1a^+/−^ mice; R. Mallarino, B. Peterson and J. Weber for experimental advice; A. Bree, E. Lievens, C. Scholes, J. Weaver and C. Xu, and for technical assistance; and S. McKnight, K. Peichel, M. Ryan, and S. Suarez for comments on earlier versions of this manuscript. Research was funded by an N.I.H. Ruth Kirschstein National Research Service Award and an N.I.H. Pathway to Independence Award to H.S.F.; an N.S.F. Graduate Research Fellowship and an N.S.F. Doctoral Dissertation Improvement Grant to E.J.-P.; a European Molecular Biology Organization Postdoctoral Fellowship and a Human Frontier Science Program Long-Term Fellowship to J.-M.L.; and an Arnold and Mabel Beckman Foundation Young Investigator Award to H.E.H. H.E.H. is an Investigator at the Howard Hughes Medical Institute.

## Author Contributions

H.S.F. and H.E.H. conceived of and designed the study; H.S.F. bred, processed and genotyped *Peromyscus*, performed QTL and fine-scale mapping, protein localization and quantification; E.J.-P. measured sperm morphology, bred and processed *Mus*; J.-M. L. designed and analysed the RNA-seq experiments; H.S.F. and H.E.H. interpreted the results and wrote the paper.

## Author Information

Sequence data from this study is available on GenBank (accession numbers KF005595 and KF005596). RNAseq data have been deposited in the GenBank/EMBL/DDBJ Sequence Read Archive under the accession codes PRJNA343919. Correspondence and requests for material should be addressed to.

